# IL-22 induced cell extrusion and IL-18-induced pyroptosis prevent and cure rotavirus infection

**DOI:** 10.1101/774984

**Authors:** Zhan Zhang, Jun Zou, Zhenda Shi, Benyue Zhang, Lucie Etienne-Mesmin, Benoit Chassaing, Andrew T. Gewirtz

**Author notes:** Corresponding and Lead Contact Author: Andrew T. Gewirtz, Ph.D., Center for Inflammation, Immunity, and Infection, Institute for Biomedical Sciences, Georgia State University, Atlanta GA 30303.

## Abstract

Administration of bacterial flagellin elicits production of TLR5-mediated IL-22 and NLRC4-mediated IL-18 that act in concert to cure and prevent rotavirus (RV) infection. This study investigated the mechanism by which these cytokines act to impede this virus. Although IL-18 and IL-22 induce each other’s expression, we found that IL-18 and IL-22 both impeded RV independently of each other and did so by distinct mechanisms, in both cases via activation of their cognate receptors in intestinal epithelial cells (IEC). IL-22 drove IEC proliferation and migration toward villus tips, which resulted in increased extrusion of highly differentiated IEC that serve as the site of RV replication. In contrast, IL-18 induced pyroptotic death of RV-infected IEC thus directly interrupting the RV replication cycle, resulting in spewing of incompetent virus into the intestinal lumen and causing a rapid drop in levels of RV-infected IEC. Together, these actions resulted in rapid and complete expulsion of RV, even in hosts with severely compromised immune systems. These results suggest that IL-18/22 might be a means of treating viral infections that preferentially target short-lived epithelial cells.

## Introduction

Rotavirus (RV) remains a scourge to humanity, causing severe distress to many and thousands of childhood deaths annually, particularly in developing countries wherein RV vaccines have only moderate efficacy [1]. RV is a double-stranded RNA virus that primarily infects intestinal epithelial cells (IEC) that line the villus tips of the ileum, resulting in severe life-threatening diarrhea in young children and moderate gastrointestinal distress in adults [2-4]. Such tropism and pathogenesis is faithfully recapitulated in RV-infected mice making the mouse model of RV useful for studying basic aspects of RV immunity and disease pathophysiology. Further, the RV mouse model may prove a useful platform for discovery of novel means to treat and prevent RV infection, especially in scenarios when adaptive immunity, which normally plays an essential role in clearing RV, is not functioning adequately. Toward this end, we previously reported that administration of bacterial flagellin rapidly cured, and/or protected against, RV infection. Such protection was independent of interferon and adaptive immunity and dependent upon generation of both toll-like receptor 5 (TLR5)-mediated IL-22 and NOD-like receptor C4 (NLRC4)-mediated IL-18, which together, resulted in prevention and/or cure of RV infection, and its associated diarrhea [5]. However, the mechanisms by which these cytokines impede RV infection remained unknown and hence was the focus of this study. Herein, we report that IL-22 acts upon IEC to drive proliferation, migration, and ultimately extrusion of infected IEC into the intestinal lumen while IL-18 drives rapid necrotic/pyroptotic death of RV-infected IEC. Together, such actions of IL-22 and IL-18 eliminate RV from the intestine independent of adaptive immunity.

## RESULTS

### IL-22 and IL-18 activate their receptors on epithelial cells to protect against rotavirus

We previously reported that systemic administration of bacterial flagellin elicits TLR5-mediated production of IL-22 and NLRC4-mediated generation of IL-18 that can act in concert to prevent or treat rotavirus (RV) and some other enteric viral infections [5]. Specifically, as shown in Figure 1A and our previous work, the chronic RV infections that developed in RV-inoculated immune-deficient C57BL/6 *Rag-1*^*-/-*^ mice were cured by combined systemic treatment with IL-18 and IL-22 while injection of either cytokine alone reduced RV loads but did not clear the virus, regardless of cytokine dose and duration of administration. While in these particular experiments, RV infection was assayed by measuring fecal RV antigens by ELISA; assay of RV genomes in the intestine yields very similar results [5]. In WT mice, while sufficiently high doses of recombinant IL-22 can, by itself, fully prevent RV infection, at lower doses exogenously administered IL-22 and IL-18 reduced the extent of RV infection, the combination of these cytokines eliminated evidence of infection (Figure 1B). The central goal of this study was to elucidate mechanisms by which these cytokines act in concert to treat and prevent RV infection.

**Fig. 1.**
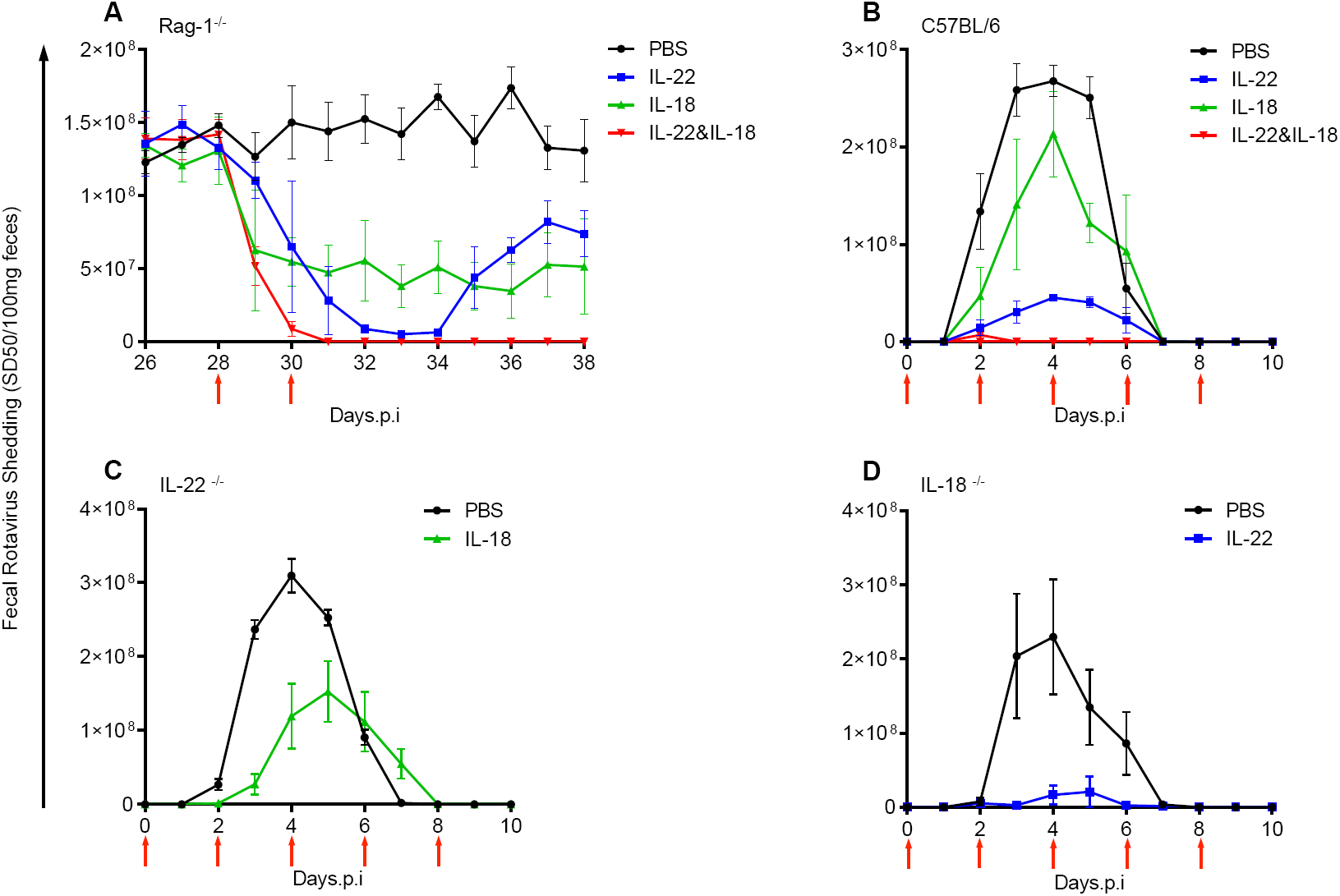
IL-22 and IL-18 elicit distinct antiviral activities against mRV invasion. (**A**) Chronically mRV-infected *Rag-1*^-/-^ mice were intraperitoneally (*i.p*) administrated with 200 µl PBS (vehicle), 10 µg IL-22, 1 µg IL-18 or 10 µg IL-22 plus 1 µg IL-18 at days 28 and 30 post inoculation as indicated with arrows. The abundance of mRV antigen was detected by enzyme-linked immunosorbent assay (ELISA), and the statistical significance of viral titers was determined by two-way analysis of variance (ANOVA) (*n*=4-5, *P<*0.0001). (**B**) 4 groups of adult C57BL/6J mice were challenged with a dose of either PBS, 2 µg IL-22, 1 µg IL-18 or both cytokines by means of intraperitoneal injection, 2 hours prior to mRV inoculation. Cytokines treatments were administered every other day from day 0 to 8 post infection. Experiment results shown among groups were significantly different (two-way ANOVA, *n*=4, *P<*0.0001). (**C** and **D**) Genetically modified mouse strains were orally inoculated with mRV. Cytokines were administered to mice 2 hours prior to inoculation, and thereafter every other day till day 8 via *i.p* injection. *IL-22*^*-/-*^ mice were treated with IL-18 (**C**), while *IL-18*^*-/-*^ mice were treated with IL-22 (**D**). The difference between mice given PBS and cytokine was statistically significant for (C) and (D) (two-way ANOVA, *n*=5-8, *P<*0.0001).

In the context of parasitic infection, both IL-18 and IL-22 promote expression of each other and loss of either impairs immunity to *Toxoplasma. gondii* [6]. Hence, we hypothesized that administration of IL-18 might impede RV primarily as a result of its previously reported ability to induce IL-22 expression. This hypothesis predicted that ability of IL-18 to protect against RV infection would be largely absent in *IL-22*^*-/-*^ mice. However, that administration of IL-18 upon RV inoculation clearly reduced the extent of RV infection in *IL-22*^*-/-*^ mice argued strongly against this hypothesis (Figure 1C). Next, we considered the converse hypothesis, namely that IL-22 might impede RV infection by elicitation of IL-18 but, analogously, observed that recombinant IL-22 markedly prevented RV infection in *IL-18*^*-/-*^ mice (Figure 1D). Thus, while IL-18 and IL-22 may well play important roles in inducing each other’s expression, our results indicate that they also each activate distinct signaling pathways that cooperate to impede RV infection.

Next, we examined the extent to which IL-18 and IL-22 acted upon the hematopoietic or non-hematopoietic compartment to impede RV infection. We used WT, *IL-18-R*^*-/-*^, and *IL-22-R*^*-/-*^ mice to generate irradiation bone marrow chimeric mice that expressed the receptors for IL-22 or IL-18 in only bone marrow-derived or radioresistant cells. Such mice were inoculated with RV, treated with recombinant IL-22 or IL-18, and RV infection monitored via measuring fecal RV antigens by ELISA. Mice that expressed the IL-22 receptor only in bone marrow-derived cells were not protected from RV infection by IL-22 (Figure 2A), whereas mice with IL-22 receptor only in radioresistant cells were strongly protected by this cytokine (Figure 2B). These results suggest that IL-22 protects mice from RV infection by acting on IEC, which is known to be populated from radioresistant stem cells and responsive to IL-22 [7]. In accord with this notion, we observed that multiple IEC cell lines are responsive to IL-22 *in vitro* via STAT3 phosphorylation although IL-22, like flagellin and IL-18, did not impact RV infection *in vitro* (Figure S1). Studies with IL-18-R chimeric mice similarly revealed that expression of this receptor in only bone marrow-derived cells conferred a modest reduction in the extent of RV infection upon IL-18 administration (Figure 2C) although the impact of IL-18 on RV infection was clearly more evident in mice that expressed IL-18-R in only radioresistant cells (Figure 2D), likely IEC. Together, these results that agonizing IL-18 and IL-22 receptors on IEC result in generation of signals that impede RV *in vivo* but not *in vitro*.

**Fig. 2.**
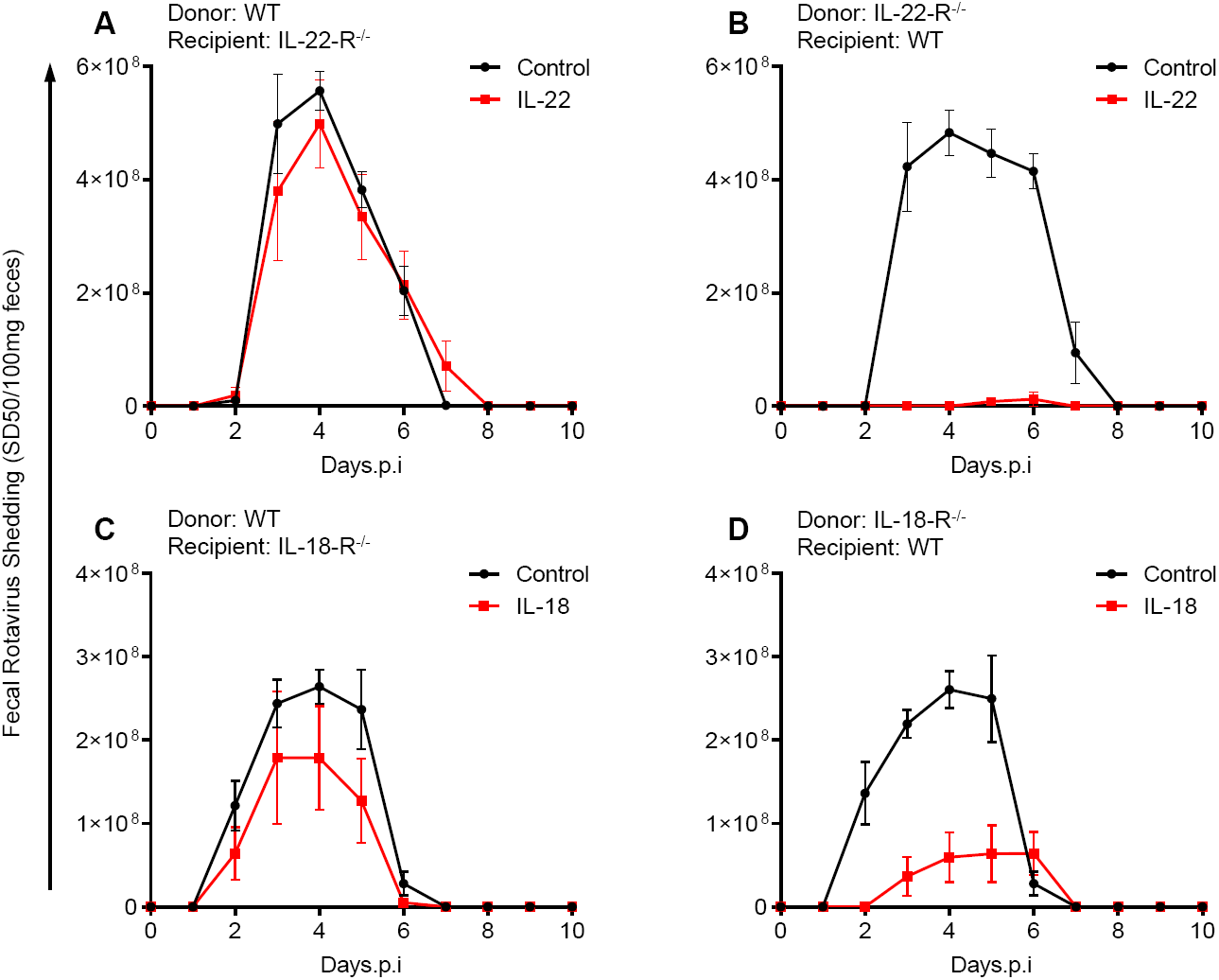
Both IL-22 and IL-18-mediated antiviral pathway requires non-hematopoietic cell compartment. Bone marrow chimeric mice from panel (**A** to **D**) were inoculated with mRV. Cytokines were given to mice 2 hours prior to inoculation, and thereafter every other day till day 8 via *i.p* injection. Feces were collected daily and assayed for mRV antigens by ELISA. Statistical evaluation was performed by two-way ANOVA. Lack of cytokines’ receptor expressed on non-hematopoietic cell compartment largely compromised the antiviral effect, as non-significant differences were discovered between PBS and cytokine-treated group (**A**) (two-way ANOVA, *n*=5-8, P=0.7715) and (**C**) (two-way ANOVA, *n*=6-8, *P<*0.0001), whereas chimeric mice groups (**B**) (two-way ANOVA, *n*=6-8, *P<*0.0001) and (**D**) (two-way ANOVA, *n*=6-8, *P<*0.0001) with consecutively expressed cytokines’ receptor on non-hematopoietic cells compartment remain their protection against mRV infection.

### IL-22 and IL-18 promotes IEC proliferation/migration

In cell culture and organoid models, IL-22 promotes IEC proliferation, migration, and stem cell regeneration [8-10], which together are thought to contribute to ability of IL-22 to promote healing in response to an array of insults including exposure to radiation and dextran sodium sulfate (DSS) *in vivo* [11-14]. In contrast to such severe injuries, RV infection is generally characterized by a lack of overt intestinal inflammation [15, 16]. Nonetheless, we envisaged that promoting IEC proliferation and/or migration, IL-22 might reduce extent of RV infection by increasing the rate of turnover of IEC, especially cells near villus tips, which is the predominant site of RV infection [2-4]. We further reasoned that, perhaps IL-18 might share such actions and thus would further increase IEC proliferation and turnover. To begin to examine these possibilities, mice were administered BrdU and treated with IL-22 and/or IL-18. Sixteen hours later, mice were euthanized and intestines subjected to fluorescence microscopy to measure rates at which IEC migrated toward villus tips, from where they are extruded into the lumen [17]. In accord with our hypothesis, administration of IL-22 approximately doubled the rate at which IEC migrated toward villus tips (Figure 3A, B). IL-18 administration also increased rate of IEC migration albeit to a lesser extent. Yet, the combination of these cytokines did not result in a faster rate of IEC migration relative to IL-22 alone. Epidermal growth factor (EGF) is known to promote IEC proliferation and migration [18, 19]. Hence, we next tested whether this cytokine might protect against RV infection. We observed EGF indeed had ability to reduce extent of RV infection (Figure 3C). Together these results support the hypothesis that promoting IEC replication and migration contributes to ability of IL-22 and IL-18 to protect against RV infection but was not able to explain the ability of these cytokines to work cooperatively toward this end.

**Fig. 3.**
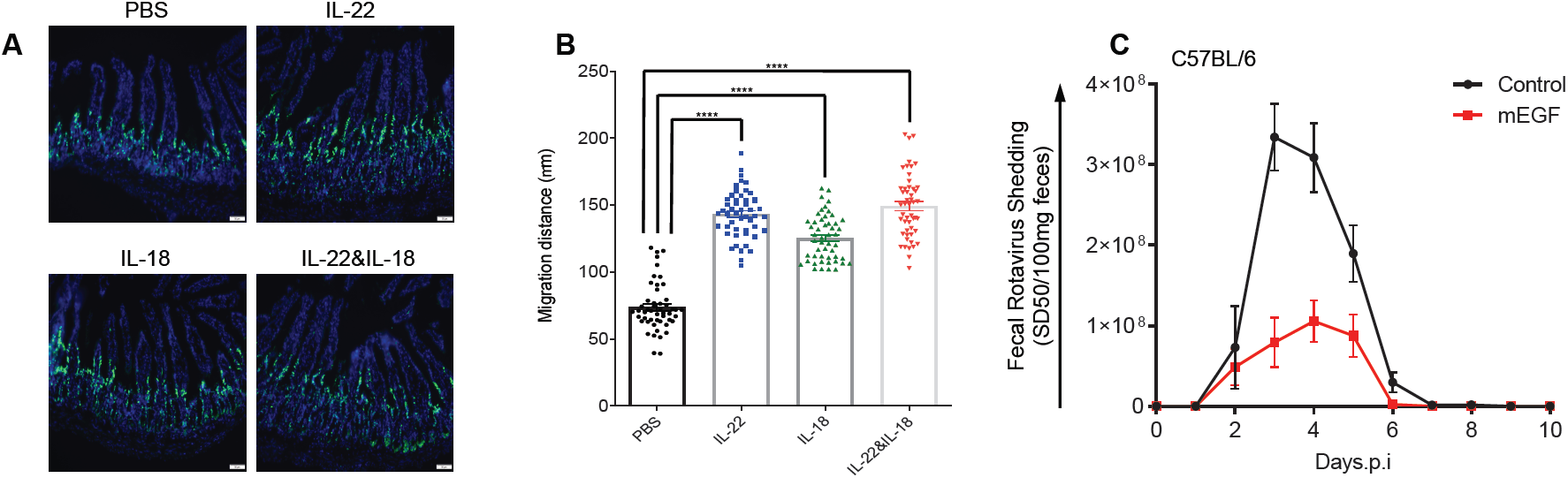
Accelerated proliferation rate and migration levels of IEC are correlated with debilitation of mRV infectivity. (**A** and **B**) Adult C57BL/6 mice were *i.p* injected with PBS, or 10 µg IL-22, 2 µg IL-18 either alone or both, following BrdU administration 1-hour post cytokine treatment. All mice were sacrificed together post 16-hour BrdU administration. (**A**) Immunohistochemistry of anti-BrdU allowed visualization of positive BrdU-labeled IEC. (**B**) Sections were scored at least from 50 villus per group of mice (n=5). Distance of the foremost migrating cells along the crypt-villus axis were measured with *ImageJ* software. Results are presented as mean ± SEM. Statistical significance was evaluated by one-way ANOVA (\*\*\*\**P<*0.0001). (**C**) Adult C57BL/6 mice were *i.p* injected with PBS, or 10 µg murine EGF 2 hours prior to mRV inoculation on day 0, and thereafter every other day from day 2 to 8 p.i. Feces were collected from both groups of mice daily, and assayed for detection of RV antigen by means of ELISA. Levels of mRV shedding are shown as mean ± SEM. The difference between PBS and mEGF-treated groups of mice were significant (two-way ANOVA, N=5, *P<*0.0001).

### IL-22 promotes extrusion of IEC into small intestinal lumen

We next considered how promoting IEC proliferation might impede RV. One seemingly likely consequence of increased IEC proliferation/migration might be increased extrusion of IEC into the lumen, which is thought to occur such that cells remain alive until extrusion is completed thus allowing the gut barrier to not be compromised [20]. Hence, we hypothesized that increased proliferation/migration induced by IL-22 and/or IL-18 treatments might result in increased extrusion of villus tip cells, which are the site of RV infection. We first investigated this hypothesis by an approach used by others [21], namely examining cross sections of H&E stained pieces of ileum for visual evidence of cell shedding, but found it difficult to distinguish IEC from other luminal contents (data not shown). Therefore, we sought to visualize such cells via a DNA stain, DAPI. While this approach suggested greater presence of IEC in the lumen of mice treated with cytokines, particularly IL-22 (Figure 4A), it was difficult to quantitate such a difference via cell counting. Hence, analogous to approaches used to quantitate gut bacteria via their 16s DNA, we sought to evaluate levels of host cells via qPCR of 18s DNA in the ileum. While the highly degradative environment of the intestine would likely degrade IEC shed into the lumen, we reasoned that since such cells are extruded in a relatively intact state, their DNA might survive long enough to enable quantitation by qPCR. Hence, as detailed in Methods, small intestinal contents were extracted and 18s DNA quantitated and expressed as number of cells per 100 mg of luminal content using known numbers of mouse epithelial cells to generate a standard curve. This approach indicated that, indeed, IL-22 treatment markedly increased the level of IEC present in the lumen (Figure 4B) indicating increased IEC shedding. IL-18 induced only a modest level of IEC shedding that appeared to be additive to the shedding induced by IL-22. A generally similar pattern was observed in the cecum (Figure 4D). In contrast, these cytokines did not impact levels of 18s DNA present in the lumen of the colon (Figure 4E), perhaps reflecting that the impact of these cytokines on IEC shedding is specific to the ileum/cecum and/or that the DNA of shed IEC is quickly degraded in the bacterial-dense colon. An even greater amount of shedding of IEC into the ileum was induced by treating mice with flagellin although 2 treatments of IL-18/22 could match this level suggesting that production of these cytokines might be sufficient to recapitulate the IEC shedding (Figure 4C) induced by flagellin. Moreover, use of *IL-22*^*-/-*^ and *IL-18*^*-/-*^ mice revealed that these cytokines, both of which are necessary for flagellin’s anti-RV action [5], were both absolutely necessary for flagellin-induced cell shedding (Figure 4F). Collectively, these results support the notion that increased extrusion of IEC, particularly in response to IL-22 might be central to this cytokine’s ability to impede RV infection but did not offer much insight into how IL-22 and IL-18 cooperate to offer stronger protection against this virus.

**Fig. 4.**
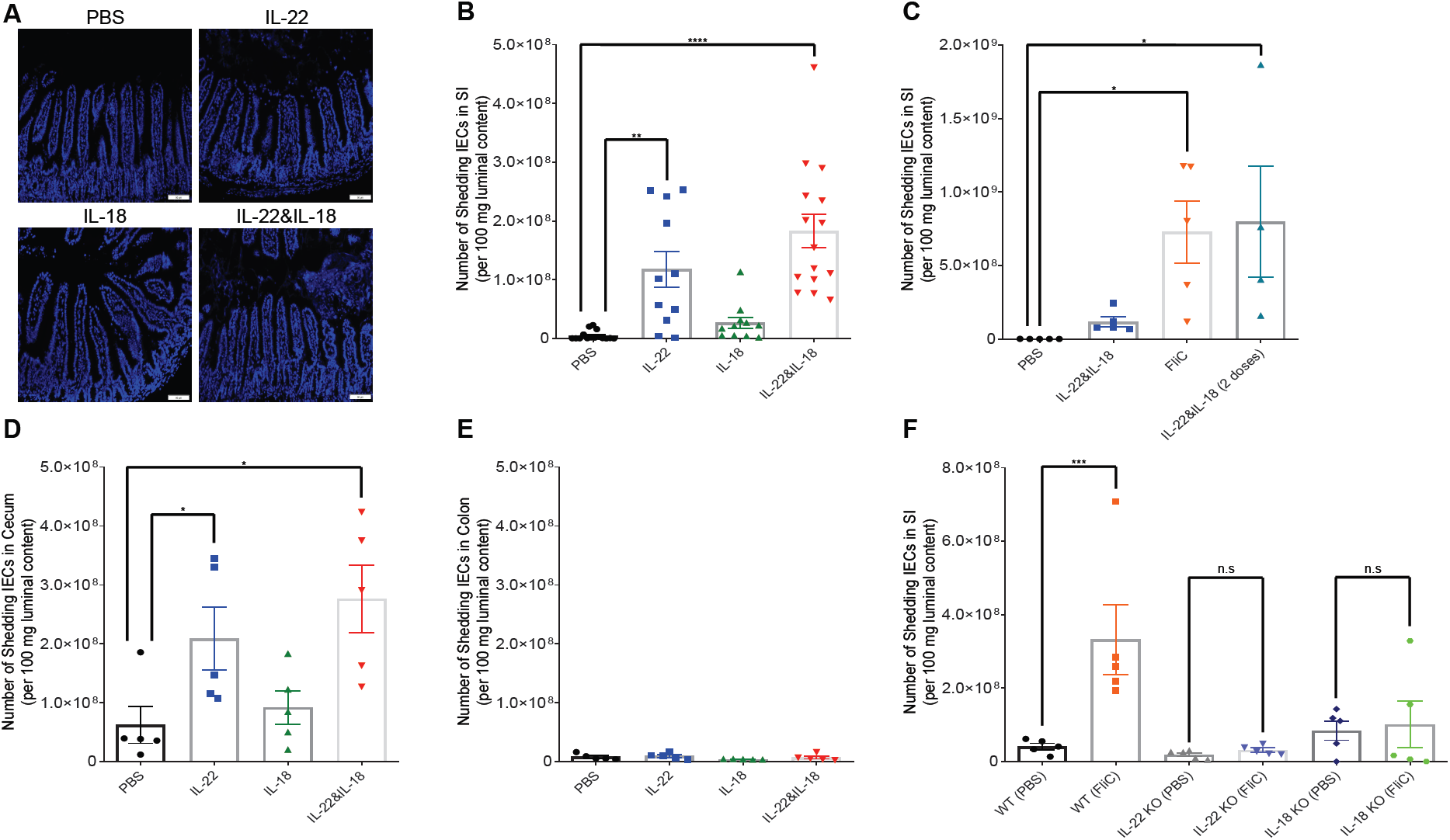
IL-22 and IL-18 mediate the elevated frequency of cell extrusion and shedding. Adult mice were *i.p* given 10 µg IL-22, 2 µg IL-18, 10 µg IL-22 plus 2 µg IL-18 or 15 µg FliC, respectively. Following 8-hour post cytokine(s) or FliC treatment, small intestines and luminal content from small intestines, cecum and colon were collected from mice. (**A**) Immunohistochemistry of C57BL/6 small intestinal sections that were counterstained with DAPI allowed visualization of shedding cells from luminal side. (**B** to **F**) Luminal content from the different regions of the gastrointestinal (GI) tract was detected for host DNA level of 18s by q-PCR. Luminal content was collected from various regions of GI tract in each panel: (**B, C** and **F**) small intestine, (**D**) cecum, (**E**) colon. (**B** to **E**) Adult C57BL/6 mice were *i.p* injected with either cytokines or FliC (administration of double doses of IL-22 plus IL-18 with 12 hours interval). Statistical significance was showed in panel (**B**), (**C**) and (**D**) (one-way ANOVA, n=5-15, \**P<*0.05, \*\**P<*0.01, *\*\*\*\*P<*0.0001), while significant difference was absent in group experiment (**E**). (**F**) The following strains of mice including C57BL/6, *IL-22*^*-/-*^, and *IL-18*^*-/-*^ were subjected to 15 µg FliC treatment. Blockade of IEC shedding rate was commitment with the ablation of IL-22 or IL-18 signaling (one-way ANOVA, n=5, *\*\*\*P<*0.001; n.s., not significant).

### IL-18 induces death of RV-infected IEC

Next, we examined how IL-22 and IL-18 might impact IEC in the absence and presence of an active RV infection. Initially, we sought to use the chronic RV infection model but the very high variance of RV levels within such animals made this approach hard to interpret (data not shown). Hence, we utilized WT mice that had been infected with RV on day 3 post-inoculation, a time approaching peak levels of RV shedding (Figure 1B). Such RV infected mice (or uninfected) mice were administered IL-22 and/or IL-18, euthanized 6 h later, and small intestinal content isolated. Like IL-18/22 administration, RV infection, by itself, upregulated IEC extrusion with a marked further increase in IEC extrusion being observed by administration of IL-18/22 to RV-infected mice (Figure 5A). This suggests that increased IEC extrusion may normally contribute to innate defense against RV [2] and that exogenously administered IL-18/22 (or flagellin) enhance this protective mechanism. Yet, like the case in uninfected mice, the promotion of IEC extrusion seemed driven by IL-22 and not IL-18 (Figure 5B).

**Fig. 5.**
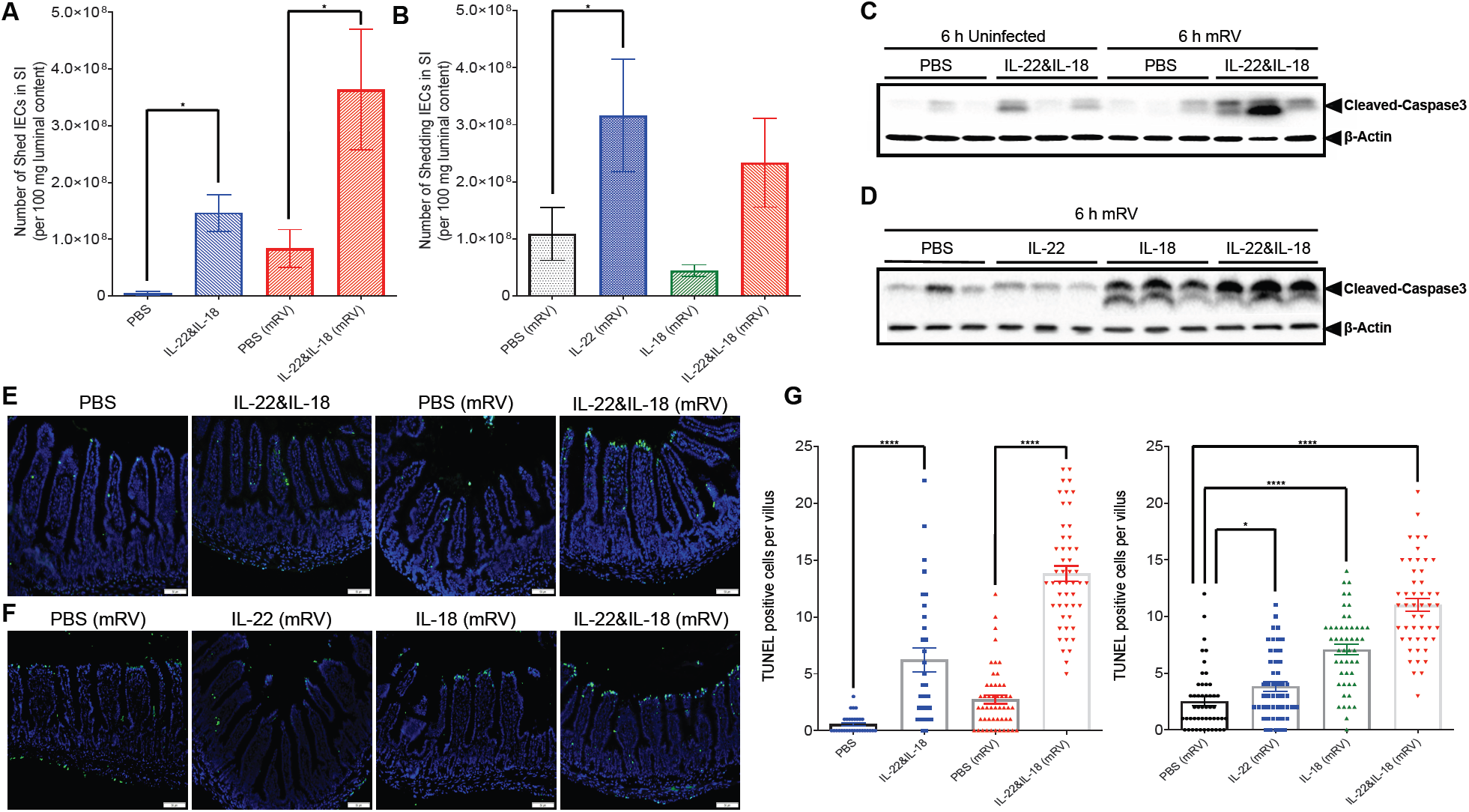
Accelerated IEC apoptosis is concomitant with the treatment of IL-18. Both infected and uninfected adult C57BL/6 mice were *i.p* given PBS, 10µg IL-22, 2 µg IL-18 or 10 µg IL-22 plus 2 µg IL-18. Administration of cytokine(s) to mRV-infected mice was on day 3 post mRV inoculation. Following 6-hour PBS or cytokines treatment, a small portion of the proximal jejunum as well as the whole luminal content from the small intestines were collected from the mice, while the rest of the small intestine was harvested to isolate the IEC. (**A** and **B**) the abundance of host DNA level of 18s from luminal content was quantified by q-PCR (one-way ANOVA, n=5, \**P<*0.05). (**C** and **D**) Whole cell lysates from IEC were analyzed by SDS-PAGE immunoblotting for detection of cleaved caspase 3. (**E** and **F**) Immunohistochemistry of TUNEL allowed visualization of apoptotic cells along the crypt-villus axis, and cell nucleus were conterstained with DAPI. (**G** and **H**) Sections were scored at least from 30 villus per group of mice, and enumerated for TUNEL-positive cells (Student’s t test, n=5, \**P<*0.05, \*\*\*\**P<*0.0001).

Next, we sought to investigate events in IEC that remained part of the small intestine at the time of increased IEC extrusion. Specifically, we sought to examine if IL-18 and/or IL-22 might impact signals associated with necrotic/pyroptotic, cell death. First, we assayed levels of cleaved caspase-3, which is known to drive such cell death pathways by SDS-PAGE immunoblot. We observed that IL-18/22 and RV, by themselves, induced modest and variable induction of Cleaved Caspase3 while these cytokines induced marked induction of Cleaved Caspase3 when administered to RV-infected mice (Figure 5C). Such induction of Caspase3 was observed in response to IL-18 but not IL-22 (Figure 5D). Quantitation of cell death by TUNEL staining also indicated that both IL-18/22 and RV by themselves resulted in an increase in TUNEL-positive cells with an approximate 2-fold further elevation being observed when IL-18/22 was administered to RV-infected mice. Yet, the localization of the TUNEL-positivity was quite striking. Specifically, IL-18/22 and RV by themselves resulted in sporadic TUNEL-positive cells throughout the villus whereas administration of IL-18/22 resulted in striking TUNEL-positivity at the villus tips, which is the primary site of RV infection (Figure 5 E-G). Analogous to the Cleaved Caspase3 activity, striking induction of TUNEL-positivity at the villus tips in RV-infected mice was seen in response to IL-18 but not IL-22. These results suggest that IL-18 might impede RV infection by causing death of RV-infected cells.

### IL-18 interrupts viral replication

Lastly, we examined the extent to which IL-22-induced IEC extrusion and IL-18-induced IEC death associated with RV reduction in the ileum at 6h and 24h following administration of these cytokines. Specifically, we measured, in both the lumen and IEC, levels of RV genomes and the ratio of +/-RV strands, which reflects levels of active replication since most + strands serve in generation of RV proteins and do not get incorporated into RV virions [22]. In accord with our previous work, we observed that in the epithelium, both IL-22 and IL-18 led to a clear reduction in both the level of RV genomes and RV replication by 6h (Figure 6 A, B). In contrast, in the small intestinal lumen, there was a marked, albeit variable, increase in the level of RV genomes and a stark increase in RV +/-strand ratios 6h following administration of IL-18, the combination of IL-18 and IL-22, but not IL-22 by itself (Figure 6C, D). By 24h, levels of RV in the lumen had dropped dramatically while the miniscule levels of remaining virus appeared to not be actively replicating (Figure 6E, F). Collectively, these results support a model wherein IL-18-induced cell death interrupts active RV replication, spewing incompletely replicated virus into the lumen while IL-22 induces IEC migration and subsequent extrusion of the mature IEC that RV targets, thus together working in concert to resolve RV infection.

**Fig. 6.**
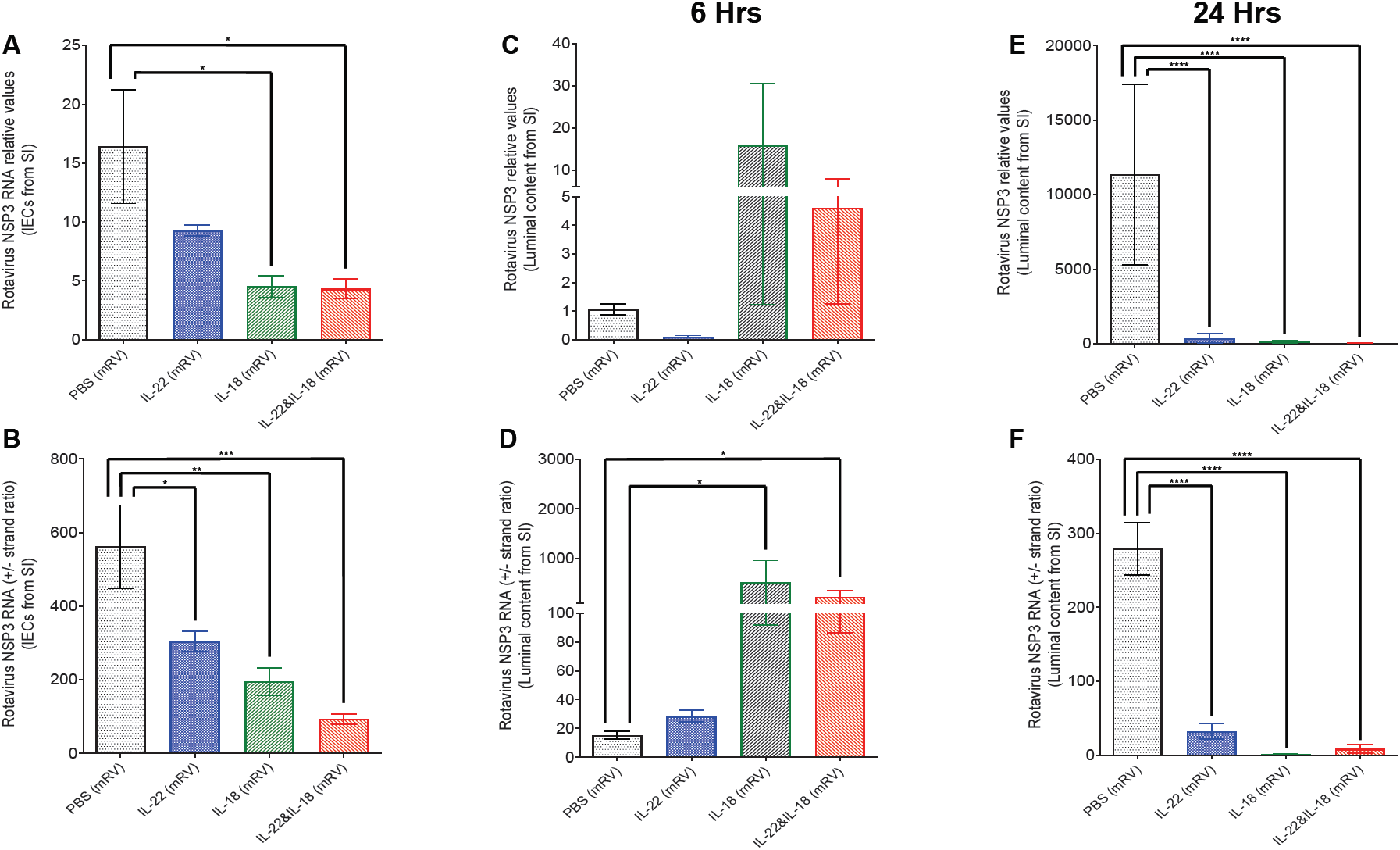
Administration of IL-18 rapidly releases the replicating virus into the luminal side. mRV-infected adult C57BL/6 mice were *i.p* injected with PBS, 10µg IL-22, 2 µg IL-18 or 10 µg IL-22 plus 2 µg IL-18 on day 3 post virus inoculation. Following 6-hour PBS or cytokines treatment, the mRV genome were extracted from the isolated small intestinal epithelial cells as well as the whole luminal content. The abundance of virus genome is reflected by NSP3 RNA levels, meanwhile the efficiency of viral replication is represented as the excess copy number of NSP3 (+) RNA strand over complimentary NSP3 (-) RNA strand. (**A** and **B**) The overall mRV genome and efficacy of virus replication in small intestinal epithelial cells. (**C** to **F**) The overall mRV genome and efficacy of virus replication in luminal content from small intestine (Student’s t test, n=5-10, \**P<*0.05, *\*\*\*\*P<*0.0001).

## Discussion

The central focus of this study was to determine the mechanism by which IL-18 and Il-22, which are elicited via bacterial flagellin, cures, and/or prevents, rotavirus infection. We initially considered the possibility that the ability of IL-18 and IL-22 to promote each other’s expression allowed them to cooperate to promote RV clearance by a common mechanism. However, we found that, irrespective of such mutual promotion, IL-18 and IL-22 both impeded RV independent of each other and did so by distinct mechanisms. Specifically, IL-22 drove intestinal epithelial cells (IEC) proliferation and migration toward villus tips, which resulted in increased extrusion of highly differentiated IEC that serve as the site of RV replication. In contrast, IL-18 induced pyroptotic death of RV-infected IEC thus directly interrupting the RV replication cycle and causing a rapid drop in levels of RV-infected IEC. We conclude that, together, these actions result rapid and complete expulsion of RV, even in hosts with severely compromised immune systems.

RV does not induce detectable increases in IL-22 expression nor does genetic deletion of IL-22 appear to markedly augment RV infection [5], thus arguing that IL-22 does not normally play a major role in clearance of this pathogen. Nonetheless, the downstream action of IL-22, particularly its promotion of IEC turnover may be shared by endogenous anti-RV host defense mechanisms. While the role of adaptive immune-independent host defense against RV is most easily appreciated in immune compromised mice wherein RV loads decline markedly from their peak levels, it may also play a role in protecting against RV even in immune competent mice. While innate host defense against RV is likely multifactorial, and may involve type III interferon [3], our observation that RV infection increases IEC extrusion, combined with previous observation that RV infection activates intestinal stem cell proliferation suggests a role for increased IEC turnover in limiting RV infection [2]. Hence, we presume that IL-22 is but one means, albeit a potent one, of activating a very basic primitive mechanism of host defense against a variety of challenges.

IEC are highly rapidly proliferating cells with average lifetimes of about 3 days [22]. Hence, the intestine must continuously eliminate vast numbers of cells. The overwhelming majority of IEC are eliminated via cell extrusion at villus tips via a process termed anoikis. A central tenet of anoikis is that cells remain alive at the time of extrusion followed by the lack of attachment to other cells resulting in induction of a programmed death process [23]. A key aspect of this process is that it permits elimination of cells without comprising gut barrier function and thus avoiding infectious and inflammation that might otherwise result therefrom. Accordingly, administration of IL-22 is associated with few adverse effects and in a variety of scenarios shows clear ability to resolve inflammation [24]. It is possible that increasing anoikis via IL-22 results in extrusion of RV-containing cells in a manner that prevents viral escape and, consequently, infection of other IEC. However, IL-22’s lack of induction of a detectable increase in luminal RV argues against this possibility. Rather, we envisage that the cell death process that follows IEC extrusion might result in destruction of RV in those cells. Additionally, and/or alternatively, we hypothesize that the accelerated IEC turnover induced by IL-22 results in villus IEC being less differentiated and thus less susceptible to RV infection. In accord with this possibility, we’ve observed that that flagellin administration resulted in an IL-22-dependent increase in CD44^+^26^-^ IEC (Figure. S2), which are known to be RV-resistant [25]. While it is difficult to discern the relative importance of IL-22’s induction of IEC extrusion versus its impact on differentiation state of villus IEC, that IL-22-induced reduction in RV levels in chronically infected *Rag-1*^*-/-*^ mice occurs over a course of several days supports a role for the latter mechanism. Use of IL-22 receptor bone marrow chimera mice demonstrated that this cytokine’s impact on RV is mediated by its direct impact on IEC [7].

In contrast to IL-22, recent work indicates induction of IL-18 plays a role in endogenous innate immunity against RV. Specifically, Zhu *et al*. demonstrated a role for RV-induced increases in IL-18, mediated by activation of the NLR9pb inflammasome, in mediating clearance of this virus in immune competent mice. Such IL-18 induction correlated with, and was necessary for, gasdermin-dependent pyroptosis, the absence of which resulted in delayed clearance of RV [26, 27]. Based on this work, we hypothesize that exogenously administered IL-18 might enhance RV-induced death of RV-infected cells and/or more generally increase IEC turnover analogous to IL-22. In accord with the latter, administration of IL-18 in the absence of RV elicited a modest increase in the number of TUNEL positive cells as well as a modest increase in IEC proliferation/migration that was not accompanied by increased IEC extrusion suggesting the increased proliferation compensated for cell death. However, such TUNEL positive cells were scattered along the villus rather than being concentrated toward the tips where RV would be located. In contrast, in RV infected mice, IL-18 led to TUNEL positive cells at the villus tips in a manner that strongly implicated pyroptosis in mediating IL-18’s anti-RV effect. These results suggest that induction of IL-18 receptor-mediated signaling by itself is not sufficient to induce cell death in villus tip epithelial cells but rather triggers death only in cells primed as a result of RV infection. The nature of such priming is not understood but could conceivable involve IEC signaling pathways, including NLR9pb, TLR3, and PKR, that are capable of recognizing RV components and/or responding to intracellular stress in general [27-29]. In this context, the ability of IL-22 to enhance IL-18-induced TUNEL positivity in RV-infected might possibly reflect an intersection of IL-22-R and IL-18-R signaling or be a manifestation of these cytokines to promote each other’s expression.

The improved understanding of the mechanism by which IL-18/22 treats RV infection reported herein should inform use of these cytokines to treat viral infection in humans. While chronic RV infections, which occur in immune compromised humans, are one potential use of IL-18/22, there are a number of chronic viral infections in need of additional therapeutic options. Our results suggest that this cytokine treatment would likely be effective for viruses that preferentially infect villus epithelial cells and perhaps epithelial cells with high turnover rates in general. In contrast, this combination of cytokines seems unlikely to impact viruses that inhabit more long-lived cells, including hematopoietic cells that are generally not responsive to IL-22. In accord with this reasoning, we’ve observed that flagellin and IL-18/22 has some efficacy against reovirus, particularly early in infection when it infects gut epithelial cells, as well as some efficacy against influenza, which initially infects lung epithelial cells, but did not show any impact on hepatitis C virus as assayed in mice engrafted with human hepatocytes (data not shown), which are thought to be long lived cells. Nor did IL-18/22 protect mice against norovirus infection, wherein the virus infects B-cells and tuft cells [30, 31]. In contrast, human norovirus is thought to primarily infect epithelial cells, particularly in immunocompromised persons who develop chronic norovirus infections [32]. Hence, we envision that chronic rotavirus and/or norovirus infections in person with immune dysfunction might be reasonable targets of IL-18/22-based therapy.

## Materials and Methods

### Mice

All mice used herein were on a C57BL/6 background and bred at Georgia State University (Atlanta, GA). Rotavirus-infected mice were housed in an animal biosafety level 2 facility under institutionally-approved animal use protocols (IACUC # 17047). WT, *Rag-1*^*-/-*^, *IL-18*^*-/-*^, *IL-18-R*^*-/-*^, *Stat3*^*flox*^, and *Villin-cre* were purchased from Jackson Laboratories. *NLRC4*^*-/-*^, *IL-22*^*-/-*^, and *IL-22-R*^*-/-*^ mice were provided by Genentech. *TLR5*^*-/-*^ and *TLR5*^*-/-*^*/NLRC4*^*-/-*^ and WT littermates were maintained as previously described [5].

### Materials

Murine Fc-IL-22 was provided by Genentech, Inc. Murine IL-18 was purchased from Sino Biological Inc (Beijing, China). Procedures for isolation of flagellin, and verification of purity, were described previously [5]. Recombinant murine epidermal growth factor (mEGF) was purchased from PEPROTECH.

### Rotavirus infection

*Acute Models:* Age- and sex-matched adult mice (8-12 weeks of age) were orally administrated with 100 µl 1.33% sodium bicarbonate (Sigma), and then inoculated with 10^5^ SD50 of murine rotavirus EC strain. Approach used to determine SD50 has been described previously (5). *Chronic model:* 5-week-old *Rag-1*^-/-^ mice were infected with mRV (same infection procedure as described in *Acute Models*). Feces were collected 3-week post rotavirus inoculation to confirm the establishment of chronic infection. *In vitro model:* Cell culture-adapted Rhesus RV was trypsin-activated (10 µg/ml trypsin in serum-free RPMI-1640 (Cellgro) at 37°C for 30 min. The basolateral side of the polarized Caco-2 cells were stimulated with cytokines, 1.5 hours prior to expose to Rhesus RV infection as previously described (5). The upper chamber of the Transwell were infected with Trypsin-pretreated RRV and allowed for adsorption at 37°C for 40 min before washed with serum-free medium (SFM). The presence of cytokines was maintained constant throughout the experiment.

### Fecal Rotavirus Antigen Detection

Fecal pellets were collected daily from individual mouse on days 0-10 post rotavirus inoculation. Samples were suspended in PBS (10% wt./vol.) and, after centrifugation, supernatants of fecal homogenates were analyzed by enzyme-linked immunosorbent assay (ELISA) after multiple serial dilutions, more detailed descriptions of experimental procedures are previously described [5].

### Generation of Bone Marrow Chimeric mice

Mice were subjected to X-ray irradiation using 8.5 Gy equivalent followed by injection of 2×10^7^ bone marrow cells administered intravenously as previously described before (1). All mice were afforded an 8-week recovery period before experimental use. For the first 2 weeks post-transfer, mice were maintained in sterile cages, and supplied with drinking water containing 2 mg/ml neomycin (Mediatech/Corning).

### Visual assessment of IEC shed into small intestinal lumen

Intestinal sections were fixed in methanol-Carnoy’s fixative solution (60% methanol, 30% chloroform, 10% glacial acetic acid) for 48 hours at 4°C. Fixed tissues were washed two times in dry methanol for 30 min each, followed by two times in absolute ethanol for 20 min each, then incubated in two baths of xylene before proceeded to paraffin embedding. 4-µm-thin sections were sliced from paraffin-embedded tissues, and placed on glass slides after floating on a water bath. The sections were dewaxed by initial incubation at 60°C for 20 min, and following two bathes in prewarmed xylene substitute solution for 10 min each. Deparaffinized sections were then hydrated in solution with decreasing concentration of ethanol (100, 95, 70, 50, and 30%) every 5 min in each bath. Last, slides were let almost dry completely, and then mounted with Prolong antifade mounting media containing DAPI before analyzed under the fluorescence microscopy.

### Immunohistochemistry for TUNEL staining

Intestinal sections were fixed in 10% buffered formalin at room temperature for 48 hours, and then embedded in paraffin. Tissues were sectioned at 4 µm thickness and IEC death was detected by TUNEL assay using the *In Situ* Cell Death Detection Kit, Fluorescein (Roche) according to the manufacturer’s instructions.

### Immunoblot Analysis for assay of Cleaved Caspase3 and Phospho-STAT3

Intestinal epithelial cells lysate (20 µg per lane) were separated by SDS-PAGE through 4%-20% Mini-PROTEAN^®^ TGX™gel (BIO-RAD), transferred to nitrocellulose membranes, and analyzed by immnoblot, as previously described (5). Briefly, isolated IEC were incubated with RIPA lysis buffer (SANTA CRUZ BIOTECHNOLOGY) for 30 min at room temperature. Subsequently, cell lysates were homogenized with pipette, and then subjected to full-speed centrifugation. The proteins bands were detected for Cleaved Caspase3, phosphor-STAT3 and anti-β-actin (Cell Signaling), and incubated with horseradish peroxidase-conjugated anti-rabbit. Immunoblotted proteins were visualized with Western Blotting Detection Reagents (GE Healthcare), and then imaged using the ChemiDoc XRS^+^ system (Bio-RAD).

### Isolation of intestinal epithelial cells

The entire small intestine was harvested from different strains of mice according to indicate experimental design, and sliced longitudinally before washed gently in PBS to remove the luminal content. Tissues were processed and maintained in 4°C at all conditions. Cleaned tissue samples were further minced into 1-2-mm^3^ pieces, and shaken in 20 ml HBSS containing 2mM EDTA and 10 mM Hepes for 30 min. An additional step of vigorous vortexing in fresh HBSS (10 mM Hepes) after EDTA incubation would facilitate cell disaggregation. Intestinal epithelial cells (IEC) were then filtered through 70-µm nylon mesh strainer (BD Biosciences), centrifuged, and resuspended in PBS.

### Antibody Staining and Flow Cytometry Analysis

Bulk leukocytes and intestinal epithelial cells isolated above were incubated with succinimidyl esters (NHS ester)-Alexa Fluor 430, which permitted determination of cell viability. Cells were then blocked by incubation with 10 µg/ml anti-CD16/anti-CD-32 (clone 2.4G2 ATCC). 20 min later, cells were stained with fluorescently conjugated antibodies: CD26-PE (clone: H194-112, eBioscience), CD44-PECy7 (clone: IM7, eBioscience), CD45-FITC (clone: 30-F11, eBioscience), CD326-APC (clone: G8.8, eBioscience). Finally, stained cells were followed by fixation with 4% formaldehyde for 10 mins before whole cell population was analyzed on a BD LSR II flow cytometer. Collected data was carried out using FlowJo.

### Quantification of IEC shedding from luminal content

Host DNA was quantitated from 100 mg of luminal content (100 mg) from small intestine by using QIAamp DNA Stool Mini kit (Qiagen), and subjected to quantitative PCR using QuantiFast SYBR Green PCR kit (Bio-Rad) in a CDX96 apparatus (Bio-Rad) with specific mouse 18S oligonucleotides primers. The sense and antisense oligonucleotides primers used were: 18s-1F: 5’-GTAACCCGTTGAACCCCATT-3’ and 18s-1R: 5’-CCATCCAATCGGTAGTAGCG-3’. PCR results were expressed as actual numbers of IEC shedding per 100 mg of luminal content, calculated using a standard curve, which was generated using two-fold serial dilutions of mouse colon carcinoma cell line MC26. DNA was extracted from each vial with known number of MC26 cells after serial dilutions, and then Real-Time quantitative PCR was performed. The cycle quantification (Cq) values (X-axis) are inversely proportional to the amount of target genes (18S) (Y-axis), and this standard-curve plot is applied to estimate the numbers of cell shedding from luminal content based on the quantity of target copies (18S) from each sample.

### Quantification of RV genomes and replication in IEC and luminal content

To extract RNA, cell pellets were homogenized with TRIzol™ (Invitrogen), and then addition of chloroform to the homogenate allowed separation between RNA (an upper aqueous layer) and DNA plus proteins (a red lower organic layer). Further, isopropanol facilitated the precipitation of RNA, and after centrifugation, the impurities from RNA were removed by washing with 75% ethanol. RNA pellet was resuspended in RNase-free water and preceded to quantitative qRT-PCR. Total RNA from luminal content is purified from RNeasy PowerMicrobiome Kit according to the manufacturer’s instructions. Then primers that target NSP3 region: EC.C (+) (5’-GTTCGTTGTGCCTCATTCG-3’ and EC.C (-) (5’-TCGGAACGTACTTCTGGAC-3’) were applied to quantify the overall viral genomes from IEC and luminal content. RV replication was as quantitated as previously described [33].

### BrdU pulse-chase labeling analysis of intestinal enterocyte migration

A pulse-chase experimental strategy by labeling intestinal enterocytes with 5-bromo-2-deoxyuridine (BrdU) was conducted to estimate the IEC migration rate along the crypt-villus axis over a defined period of time. Briefly, 8-week-old mice were intraperitoneally injected with either PBS or cytokine(s) (IL-22 and/or IL-18) 1 hour prior to the BrdU *i.p.* treatment (50 µg per mg of mice body weight). After 16 hours, mice were euthanized, and the segment of the jejunum were resected, immediately embedded in OCT and then proceeded to tissue sectioning. 4 µm tissue sections were firstly fixed in 4% formaldehyde for 30 min at room temperature, and then washed 3 times in PBS. DNA denaturation was performed by incubating the sections in prewarmed 1.5 N HCl for 30 min at 37 °C. Then the acid was neutralized by rinsing the sections 3 times in PBS. Before BrdU immunostaining, sections were blocked with rabbit serum (BioGenex, Fremont, CA) for 1 h at room temperature, then incubated with anti-BrdU (Abcam) 2 hours at 37 °C, and counterstained with 4’, 6-diamidino-2-phenylindole (DAPI). The BrdU-labeled cells were visualized under the fluorescent microscope.

### Quantification and statistical analysis

Significance was determined using the one-way analysis of variance (ANOVA), or student’s t-test (GraphPad Prism software, version 6.04). Differences were noted as significant \**P*<0.05, \*\**P*<0.01, \*\*\**P*<0.001, \*\*\*\**P*<0.0001.

## Abbreviations

RV: Rotavirus;
IEC: intestinal epithelial cells;
SI: small intestine

---

**Fig. S1.**
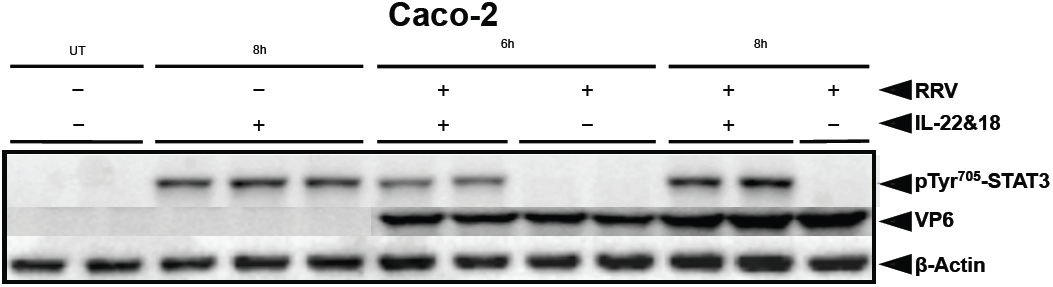
IL-22 and IL-18 couldn’t alter RV infection in cultured IEC. The basolateral side of the polarized Caco-2 cells were stimulated with 0.5 µg/ml IL-22 plus 0.25 µg/ml IL-18, 1.5 hours prior to expose to Rhesus Rotavirus (RRV) infection. Trypsin-pretreated RRV was added to the upper chamber of the Transwell plates and allowed for adsorption at 37°C for 40 min before washed with serum-free medium (SFM). The presence of IL-22 and IL-18 were maintained constant throughout the experiment. Cell lysates were collected at indicated time points and analyzed for the virus antigen VP6 and phosphorylation of STAT3. Administration of IL-22 and IL-18 successfully induced phosphorylation of STAT3, however, cytokine treatment didn’t alleviate RRV infection (Student’s t test, n=5, \**P<*0.05).

**Fig. S2.**
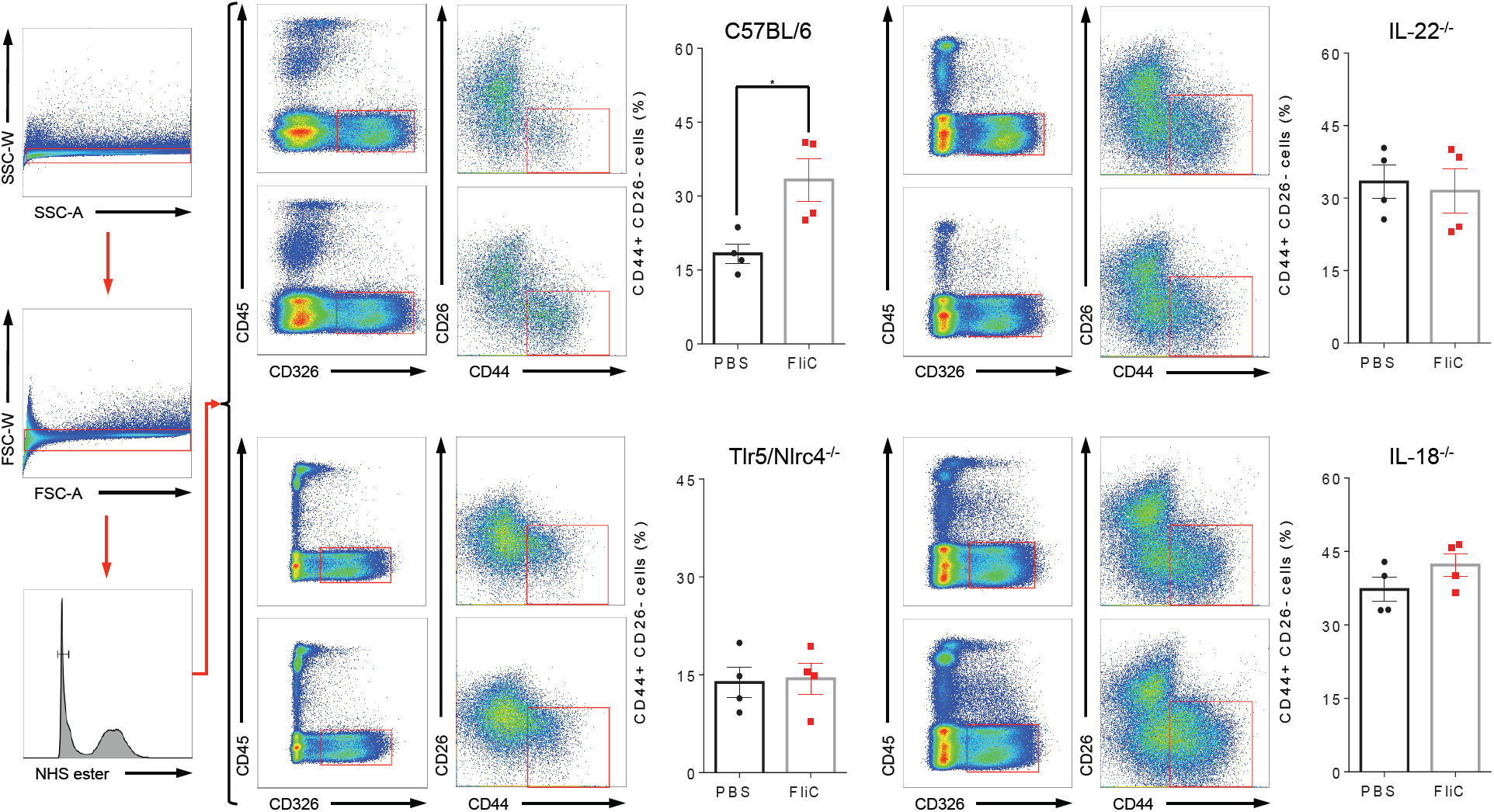
Flagellin-mediated changes of cell subpopulations along intestinal villus-crypt axis. (A to D) C57BL/6J, *TLR5*^*-/-*^*/NLRC4*^*-/-*^, *IL-22*^*-/-*^, and *IL-18*^*-/-*^ mice were treated with PBS ± flagellin (20 µg) via intraperitoneal injection. Following 24-hour PBS or flagellin administration, IEC were isolated from the small intestines. Debris and doublets were excluded by sequential gating on SSC-width vs. SSC-area, followed by FSC-width vs. FSC-area. Dead cells were excluded from alive ones based on succinimidyl esters (NHS ester)-Alexa Fluor 430. The isolated IEC (CD326^+^CD45^-^) were separated for CD26, a marker enriched in cell subpopulation that are susceptible to rotavirus infection, and CD44, is highly expressed in cells that are resistant to rotavirus infection. (A) Scatter plots using CD26 and CD45 to quantitate the percentage of IEC subsets (CD326^+^CD45^-^) in each condition from different mice strains. The difference of IEC (CD44^+^CD26^-^) subsets between the PBS and flagellin groups was statistically significant in WT C57BL/6j, while nonsignificant in *TLR5*^*-/-*^*/NLRC4*^*-/-*^, *IL-22*^*-/-*^ and *IL-18*^*-/-*^ groups (Student’s t test, n=5-10, \**P<*0.05).

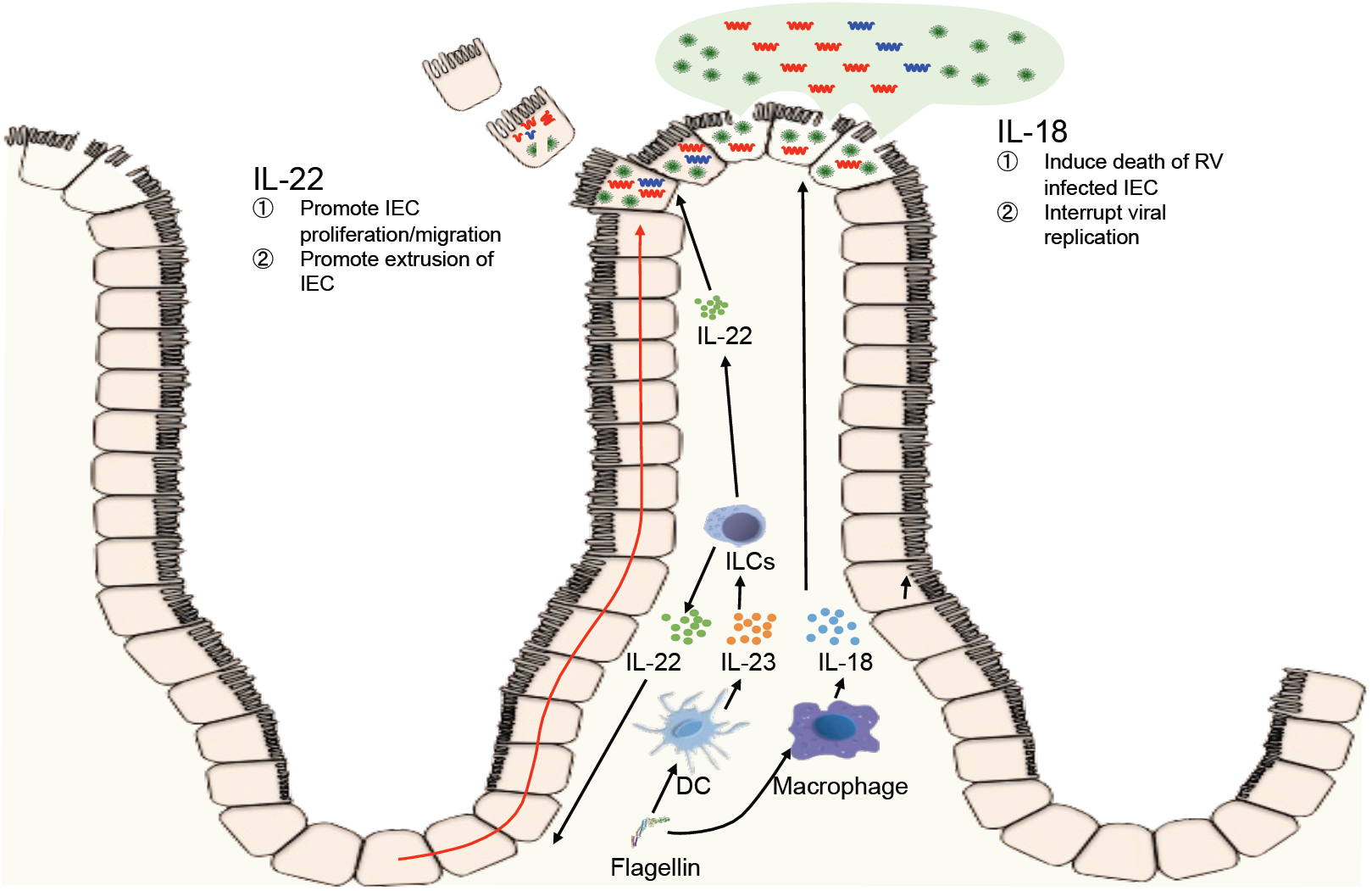

